# Spectral tuning of proton-pumping rhodopsins underscores distinct light adaptations in polar diatoms

**DOI:** 10.64898/2026.11.24.768933

**Authors:** Brittany N. Zepernick, Sarah M. Andrew, Meredith G. Meyer, Emily E. Chase, Gavin Madgett, James L. Pinckney, Sven A. Kranz, Laura Gomez-Consarnau, William G. Sunda, Adrian Marchetti

**Affiliations:** Department of Earth, Marine and Environmental Sciences, University of North Carolina at Chapel Hill, Chapel Hill, NC, USA; Department of Biology & Marine Biology, University of North Carolina Wilmington, Wilmington, NC USA; Department of Biology, University of Winnipeg, Winnipeg, MB, CA; Department of Biological Sciences and School of the Earth, Ocean and Environment, University of South Carolina, Columbia, SC, USA; Department of Biosciences, Rice University, Houston, Texas, USA; Mediterranean Institute for Advanced Studies (IMEDEA), Consejo Superior de Investigaciones Cientificas-Universitat de les Illes Balears (CSIC-UIB), Esporles, Spain

**Keywords:** Southern Ocean, Climate Change, Iron Limitation, Light Spectra, Xanthorhodopsin

## Abstract

Resource limitation constrains primary production across vast regions of the global ocean. Diatoms often persist under these conditions, despite their relatively large size and nutrient quotas compared to other phytoplankton phyla. This trend is particularly pronounced in the variable light and iron-depleted waters of the Southern Ocean. Proton-pumping rhodopsins (PPRs) have emerged as an important adaptation underlying this success, allowing diatoms to energetically supplement chlorophyll-based photosynthesis with retinal-based phototrophy, particularly under iron stress. While PPRs are increasingly recognized as a driver of diatom resilience under iron depletion, the extent to which coinciding environmental factors – such as light – shape this adaptation remains unresolved. We performed an *in vitro* study to investigate how variations in iron availability and light spectra synergistically shape PPR physiology in two ecologically relevant Southern Ocean diatoms: *Pseudo-nitzschia subcurvata* and *Synedra hyperborea*. Integrative biochemical and bioinformatic approaches identified two distinct evolutionary strategies for rhodopsin-mediated light acquisition – while *P. subcurvata* employed a specialist approach with two PPRs utilized under different wavelengths of light (green v. blue/red), *S. hyperborea* adopted a generalist approach with expression of a single PPR under both light treatments. Structural analyses of PPRs revealed hallmarks consistent with xanthorhodopsins, while coordinated transcriptional and pigment analyses suggest the blue-absorbing xanthophyll diadinoxanthin functions as a secondary light-harvesting chromophore. Collectively, these findings expand the ecological light niche of diatom rhodopsins, reveal previously unrecognized diversity in eukaryotic rhodopsin strategies, and suggest spectral tuning of proton pumping rhodopsins will enhance diatom fitness within the climatically altered light climate of the Southern Ocean.

**Significance Statement:** Proton-pumping rhodopsins enable diatoms to harvest light and generate energy independent of photosynthesis within the iron-depleted polar oceans. We report some diatoms possess two PPRs transcriptionally tuned to distinct light spectra and employ disparate strategies to exploit variable light environments under iron limitation. While *Pseudo-nitzschia subcurvata* transcriptionally tunes two PPRs to different light spectra, *Synedra hyperborea* expresses one PPR across multiple spectra. Contrary to expectations, PPR expression was highest under blue/red rather than green light – with analyses suggesting blue-absorbing diadinoxanthin functions as a secondary light-harvesting chromophore. These findings expand the ecological light niche of eukaryotic PPRs, reveal unrecognized diversity in rhodopsin-based energy acquisition, and improve our understanding of how carbon-sequestering phytoplankton will respond to future changes in light regimes.

## Introduction

Scarcity of resources such as iron, nitrogen and light constrains phytoplankton primary production across the global ocean (1, 2), and these limiting resources vary across spatiotemporal gradients (3–5). Iron limitation constrains primary production across one-third of the global ocean, with the Southern Ocean (SO) comprising the largest of these regions (1, 6). Despite this chronic iron depletion, the SO exerts a disproportionate influence on the global carbon cycle (7), with seasonally dominant diatoms mediating carbon flow within the dynamic water column (8, 9). Variations in light intensity and spectra further constrain (and often co-limit) this integral assemblage – with changes in stratification, seasonal light cycles and ice cover synergistically shaping the light climate (10–12). Consequently, understanding how polar diatoms respond to environmental variation is critical for predicting future ocean productivity and carbon sequestration. This need is increasingly urgent in the context of climate change – which is projected to drive changes in iron (13–15) and light (16, 17) throughout the water column. Yet, despite the certainty of this climatically altered future, the mechanisms posed to enhance diatom fitness under these conditions remain unresolved.

Proton-pumping rhodopsins (PPRs) have emerged as an important phototrophic mechanism enabling diatoms to persist in the dynamic water column. PPRs use the pigment retinal to harvest light and generate proton gradients which could drive ATP synthesis and active membrane transport – with PPR phototrophy suggested to be capable of capturing as much light energy as chlorophyll *a*-based photosynthesis under certain environmental conditions (18). Initially identified in halophilic archaea (19) and later marine bacteria (20, 21), PPRs were only recently discovered in marine diatoms (22). Subsequent studies revealed PPRs are particularly widespread among SO diatoms, expressed under iron limitation, and linked to increases in growth rate and biomass accumulation (23–25). In total, these findings coined the “iron limitation light switch” whereby diatoms may supplement photosynthesis with PPR phototrophy under iron stress – leveraging the reduced iron requirement for PPR maintenance (26).

While PPRs are increasingly recognized as a driver of diatom resilience under iron stress, the extent to which additional environmental factors shape this adaptation remains unclear. Chlorophyll-based photosynthesis and retinal-based PPR phototrophy are predicted to be favored under contrasting conditions beyond iron availability (27), and these two methods of phototrophy are posed to enable diatoms to exploit distinct light niches. The absorption maxima of chlorophyll *a* (Chl *a*) is constrained to blue (450-490 nm *λ*) and red (620-750 nm *λ*) wavelengths, whereas retinal primarily absorbs green wavelengths (495-570 nm *λ*) (24, 28). Thus, it has been hypothesized that PPR phototrophy would be selected for by green light under low iron availability (27). In support of this hypothesis, studies have confirmed select polar diatom PPRs possess the prokaryotic hallmark of green-light spectral tuning – a leucine residue at position 105 (23, 24, 29, 30) and demonstrated the putative PPRs of *Fragilariopsis cylindrus* and *Pseudo-nitzschia granii* exhibit maximum absorbance under green light (24, 28). Yet, beyond these reports, the extent to which light spectra shapes PPR function in polar diatoms remains unresolved. Given the strong spatiotemporal and seasonal variability in SO light regimes and their anticipated alteration under climate change (31), deducing how light spectra shapes rhodopsin phototrophy in polar diatoms is of pressing importance.

We performed an *in vitro* investigation to elucidate how light spectra modulates PPR utilization under low iron availability within *Pseudo-nitzschia subcurvata* and *Synedra hyperborea*, two SO diatoms previously confirmed to encode a PPR protein (23). Integrating complementary biochemical and transcriptional approaches, we report two mechanisms of PPR-mediated light adaptation: A specialist approach by which two distinct PPRs are transcriptionally tuned to different light qualities in *P. subcurvata*, and a generalist approach by which one PPR is transcribed under both light qualities in *S. hyperborea*. We further provide evidence suggesting blue-light absorption may be accomplished by a carotenoid-binding antenna consistent with xanthorhodopsin and hypothesize diadinoxanthin may serve as the chromophore used to access this spectra. Subsequently, we propose spectrally tuned PPRs will enhance diatom fitness within the climatically altered SO, where light spectra serve to be shifted by compounding stressors. Cumulatively, these findings expand the ecological light niche of diatom rhodopsins, reveal previously unrecognized diversity in eukaryotic rhodopsin strategies, and suggest spectral tuning of proton pumping rhodopsins will enhance diatom fitness within the climatically altered SO.

## Results

### Iron availability largely shapes P. subcurvata and S. hyperborea physiology

Iron exhibited a profound effect on diatom physiology, with light spectra exerting a lesser effect. Growth responses differed between the two SO diatoms under the experimental treatments. *P. subcurvata* growth rates did not significantly vary with light spectra or iron availability (*p* > 0.273), though rates were consistently higher under iron-replete conditions (Figure 1A). In contrast, *S. hyperborea* growth was jointly constrained by iron and light. Under green light, growth rates were ∼50% higher in iron-replete vs. iron-limiting treatments (*p* = 0.021), whereas iron had no significant effect within blue/red light treatments (*p* = 0.178). Notably, growth under iron limitation was ∼50% higher in blue/red vs. green light (*p* = 0.020) (Figure 1B).

**Figure 1.**
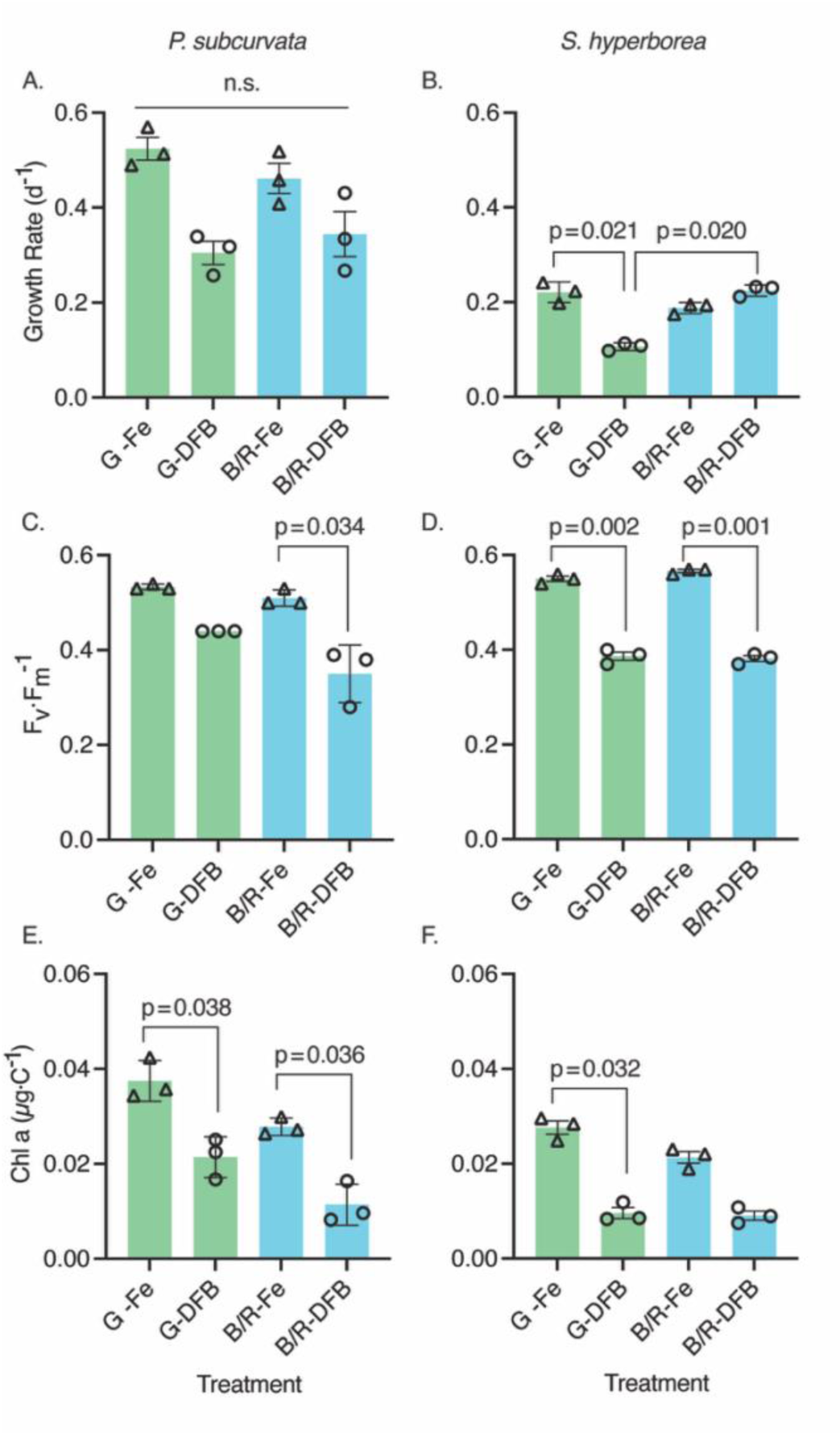
SO diatom physiology under varying iron availability and light spectra. (A) Growth rates (d^-1^) of *P. subcurvata*. (B) Growth rates (d^-1^) of *S. hyperborea*. (C) Quantum efficiency of PSII (Fv•Fm^-1^) in *P. subcurvata*. (D) Quantum efficiency of PSII (Fv•Fm^-1^) in *S. hyperborea*. (E) Chl *a* normalized to cellular carbon in *P. subcurvata* (µg Chl *a* / µg C^-1^; µg C^-1^). (F) Chl *a* normalized to cellular carbon in *S. hyperborea* (µg Chl *a* / µg C^-1^; µg C^-1^). High iron (Fe) and low iron (DFB) replicates are shown as open triangles and open circles, respectively. Green-light treatments (G) are shown in green and blue/red-light treatments (B/R) in blue. Bars indicate treatment means and black error bars denote standard error of the mean (SEM). Significant effects of light spectra or iron treatment (p < 0.05) are indicated. Here, n.s = not significant.

Photosynthetic physiology was primarily determined by iron status in both species. Maximum photochemical efficiency of photosystem II (PSII; Fv•Fm^-1^) significantly declined under low iron conditions in both light treatments (*p* < 0.034) (Figure 1C, D), though results fell short of significance in green light treatments of *P. subcurvata* (*p* = 0.097). Cellular chlorophyll *a* was significantly reduced under low iron treatments in both species (*p* < 0.038) (Figure 1E, F), with *S. hyperborea* blue/red light treatments falling short of significance (*p* = 0.067). Cellular stoichiometry suggested iron stress, with C:N ratios increased under low iron conditions in both SO species across light treatments (Supplemental Figure 1A, B). Collectively, iron availability was the primary driver of variations in growth, photosynthetic efficiency and cellular stoichiometry within both polar diatoms.

### Light spectra modulate P. subcurvata photopigment content under low-iron availability

Light constrained photopigment concentrations within *P. subcurvata* cells under low iron availability. While Chl c_3_ was only detected under green light treatments (Supplemental Figure 2A), concentrations of Chl c_1_c_2_ and fucoxanthin were significantly higher in green light across low iron treatments vs. blue/red light (p < 0.029), with β-carotene falling short of significance (*p* = 0.101) (Supplemental Figure 2B, C, D). Concentrations of diadinoxanthin (DD) and diatoxanthin (DT), which are reversibly interconverted in the diadinoxanthin cycle (DD-DT cycle) (32–34), were highest in blue/red light treatments compared to green light treatments (Supplemental Figure 2E, F). In contrast, iron was the primary determinant of photopigment concentrations across *S. hyperborea* samples irrespective of light treatment (Supplemental Figure 3A-E). Light only exerted a clear effect on cellular concentrations of DT – which were highest under blue/red light within high iron conditions (Supplemental Figure 2F). Overall, the xanthophylls DD and DT were the only photopigments that increased in the blue/red light treatments compared to green light within both SO diatoms.

### Light constrains P. subcurvata expression while iron constrains S. hyperborea expression

Whole transcriptomes clustered by light spectra across *P. subcurvata* replicates (stress = 0.0391) (Figure 2A) and iron availability across *S. hyperborea* replicates (stress = 0.0536) (Figure 2B). Consistent with these patterns, light spectra strongly shaped differential gene expression under low iron availability in *P. subcurvata,* with ∼51% of genes differentially expressed in blue/red vs. green light (*p*_adj_ < 0.05, abs Log_2_FC >1) (Figure 2C). Conversely, iron availability was the primary driver of differential expression across *S. hyperborea* libraries, with just 6% of genes differentially expressed as a function of light spectra amongst iron-limited replicates (Figure 2D). Cumulatively, SO diatoms exhibited distinct transcriptional responses: Light spectra primarily shaped *P. subcurvata* expression whereas *S. hyperborea* expression was largely influenced by iron availability.

**Figure 2.**
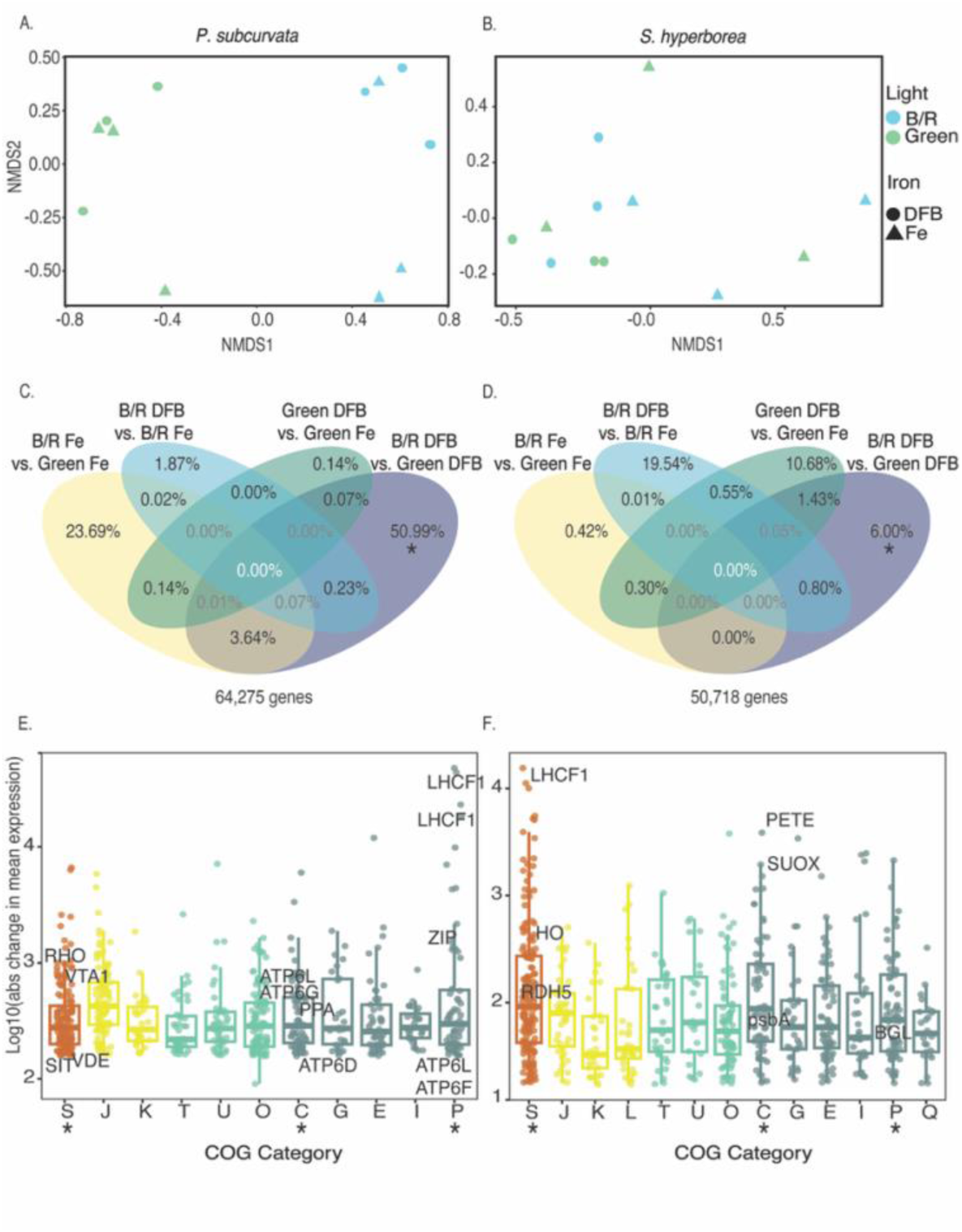
Transcriptional responses of SO diatoms to varying iron availability and light spectra. (A) nMDS of *P. subcurvata* expression (stress = 0.0391). (B) nMDS of *S. hyperborea* expression (stress = 0.0536). High iron (Fe) and low iron (DFB) replicates are shown as solid triangles and solid circles, respectively; green-light treatments (G) are shown in green and blue/red-light treatments (B/R) in blue. (C) Differentially expressed *P. subcurvata* genes. (D) Differentially expressed *S. hyperborea* genes. Venn diagram sectors represent a pairwise comparison, with the percentage of significantly differentially expressed genes indicated; overlaps denote shared genes among comparisons. The primary comparison of interest (low iron, blue/red light vs. green light) is in purple and marked (*). (E) SIMPER analysis of the top *P. subcurvata* genes contributing to dissimilarity between low iron blue/red and green light treatments. (F) SIMPER analysis of the top *S. hyperborea* genes contributing to dissimilarity between low iron blue/red and green light treatments. Log-transformed changes in absolute mean expression are grouped by clusters of orthologous groups (COGs): unknown function, S (orange); information storage and processing, J/K/L (yellow); cellular processes and signaling, T/U/O (teal); and metabolism, C/G/E/I/P/Q (dark blue). Genes of interest are labeled, and COGs of interest are marked (*).

To identify genes driving differential expression between our comparisons of interest (blue/red vs. green light qualities under low iron availability), Similarity Percentage (SIMPER) analyses were performed with the top 1,000 genes driving dissimilarity classified into clusters of orthologous groups (COGs) (Figure 2E, F). In *P. subcurvata*, genes encoding for PSII proteins drove differential expression, including psbA (gene_147945_3), psbO (gene_103002_2), psbU (gene_103002_2) and light-harvesting complex f-like genes (Lhcf1_x) (Supplemental Table 1S). Notably, genes previously associated with PPRs were strong contributors to differential expression (27), including a putative second rhodopsin-like gene (gene_140558; RHO), inorganic pyrophosphatase (gene_155322_1; PPA), and various vacuolar-ATPases (ATP6_x) (Supplemental Table 1S). Genes involved in xanthophyll biosynthesis, including violaxanthin de-epoxidase (gene_125761_4; VDE), also contributed strongly to differential expression between light treatments under low-iron conditions. In contrast, PPR-associated genes did not contribute to differential expression amongst low-iron *S. hyperborea* samples as a function of light beyond retinal dehydrogenase (gene_100515_5; RDH5). Together, these results indicate light-dependent transcriptional responses under low iron availabilities are driven in part by PPR phototrophy genes in *P. subcurvata*, whereas such patterns are less pronounced in *S. hyperborea*.

### P. subcurvata encodes two PPRs transcriptionally tuned to distinct light spectra

*P. subcurvata* was previously confirmed to possess a PPR by Andrew et al. (2023), which was found to be localized to the vacuole. This PPR was identified within this study’s coassembly (Ps_PPR1; gene_128084_2), with the 260 amino acid residues present at 100% identity. Two-dimensional structural predictions are consistent with features of rhodopsins, including seven transmembrane domains, the retinal chromophore and retinal binding pocket (23, 24) (Figure 3A). Gene expression of Ps_PPR1 was highest under green light whereas expression was largely undetectable under blue/red light (Figure 3B). Notably, expression was approximately three times higher in the low-iron treatment under green light compared to high-iron treatments under green light. Subsequent three-dimensional predictions indicated the presence of a fenestration: An opening characteristic of xanthorhodopsin (XR) which exposes the retinal-binding pocket to permit binding of a carotenoid antenna with a secondary chromophore (Figure 3C) (35–37). In total, Ps_PPR_1 was expressed highest under green light and low iron availability.

**Figure 3:**
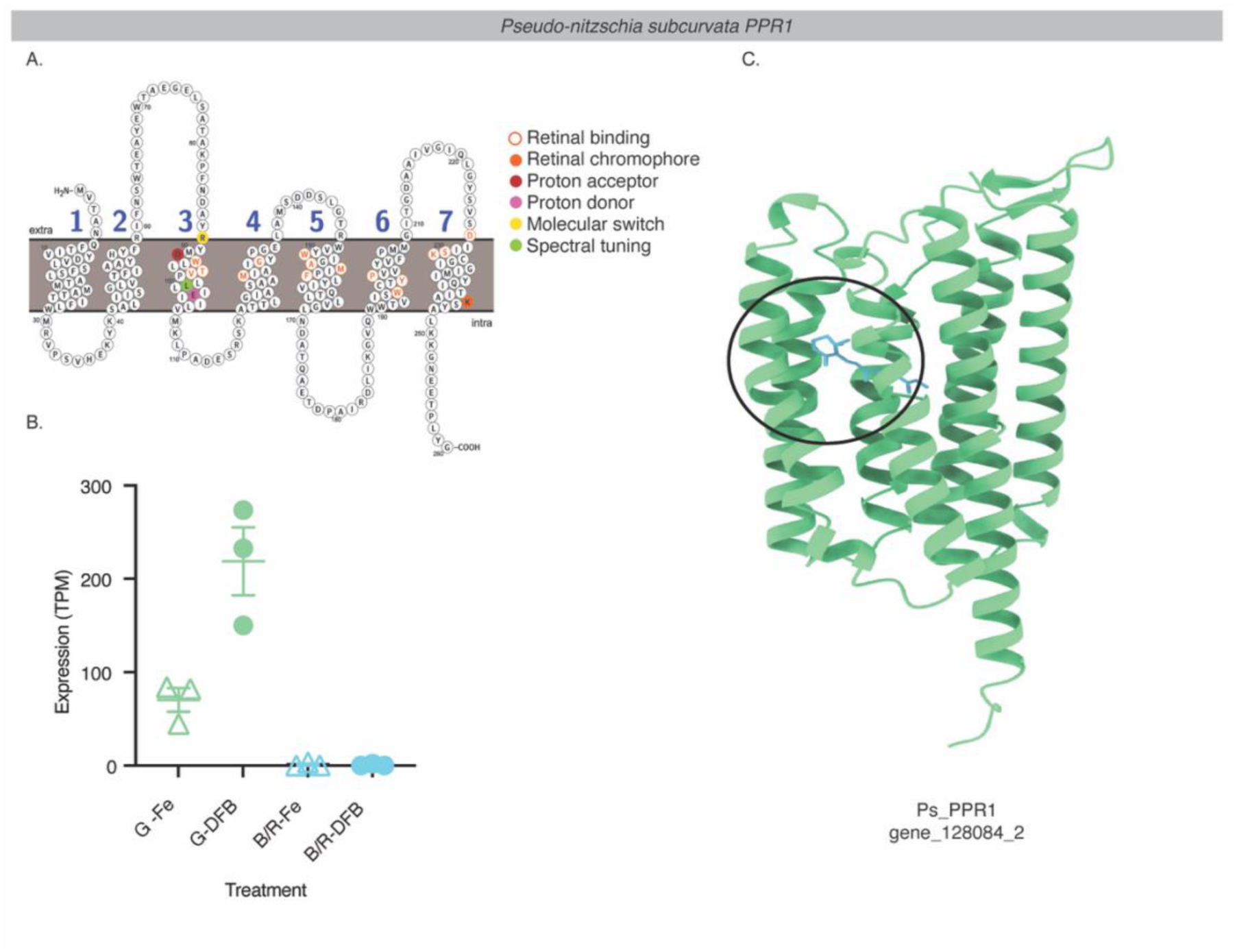
Protein structure and gene expression of green-tuned PPR of *P. subcurvata* (Ps_PPR1). (A) Two-dimensional topography of *P. subcurvata* PPR reported previously (Ps_PPR1; (gene_128084_2) (23). (B) Normalized expression (transcripts per million; TPM) of Ps_PPR1 under varying iron conditions and light spectra. High iron (Fe) and low iron (DFB) replicates are shown as open triangles and solid circles, respectively; green-light treatments (G) are shown in green and blue/red-light treatments (B/R) in blue. (C) Predicted three-dimensional structure of Ps_PPR1. The retinal β-ionone ring position within the fenestration zone is circled.

A novel, second PPR was recovered in this study which exhibited 31 discrete amino acid substitutions spanning the 261 residues (Ps_PPR2: gene_140558_3) (Figure 4A). Approximately 30% of substitutions were concentrated within (or adjacent to) the seventh transmembrane domain (residues 227-248) which contains the retinal chromophore and binding pocket. The location of these substitutions are suggested to interfere with prior immunofluorescence queries of *P. subcurvata* which targeted the C-terminus of the PPR protein (23), and we note earlier transcriptional screenings under white light were performed using a separate workflow on unassembled contigs. Notably, expression of Ps_PPR2 was approximately seven times higher in the low-iron treatment under blue/red light compared to high-iron treatments. (Figure 4B). Three-dimensional predictions indicated Ps_PP2 also possesses a fenestration characteristic of an XR (Figure 4C). In total, Ps_PPR_2 is expressed highest under blue/red light and low iron availability. More broadly, *P. subcurvata* encodes two PPRs transcriptionally tuned to distinct light qualities under low iron conditions.

**Figure 4:**
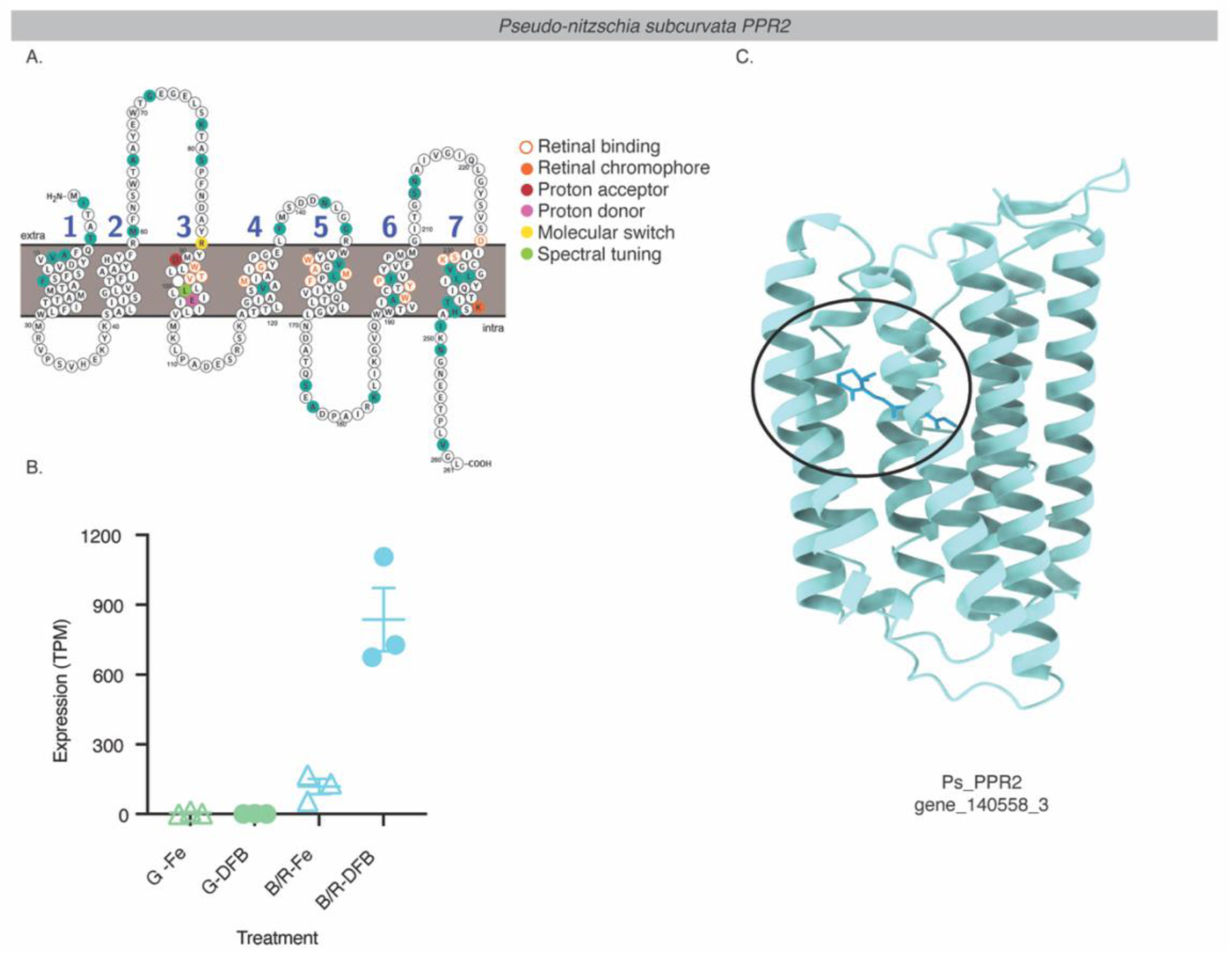
Protein structure and gene expression of novel blue/red-tuned PPR of *P. subcurvata* (Ps_PPR2). (A) Two-dimensional topography of the novel *P. subcurvata* PPR reported in this study (Ps_PPR2; k123_140558_3). Amino acid substitutions are in teal. (B) Normalized expression (transcripts per million; TPM) of Ps_PPR2 under varying iron conditions and light spectra. High iron (Fe) and low iron (DFB) replicates are shown as open triangles and solid circles, respectively; green-light treatments (G) are shown in green and blue/red-light treatments (B/R) in blue. (C) Predicted three-dimensional structure of Ps_PPR2. The retinal β-ionone ring position within the fenestration zone is circled.

### S. hyperborea encodes one PPR transcriptionally tuned to both light spectra

*S. hyperborea* was previously confirmed to possess a PPR (23), which was recovered from the coassembly at 100% identity (Sh_PPR; gene_94996_1). Consistent with PPRs recovered in *P. subcurvata*, two-dimensional structural predictions confirmed Sh_PPR contains the hallmark features of rhodopsins (Figure 5A). Yet, in contrast to trends observed in *P. subcurvata*, Sh_PPR was expressed under both light treatments. Expression was highest under low iron conditions compared to high iron conditions (Figure 5B), and transcription ∼50% higher in blue/red light compared to green light. Three-dimensional predictions indicated Sh_PPR also possesses a fenestration characteristic of an XR (Figure 5C). In total, these results suggest *S. hyperborea* encodes one PPR transcriptionally tuned to both light spectra under low iron availability.

**Figure 5:**
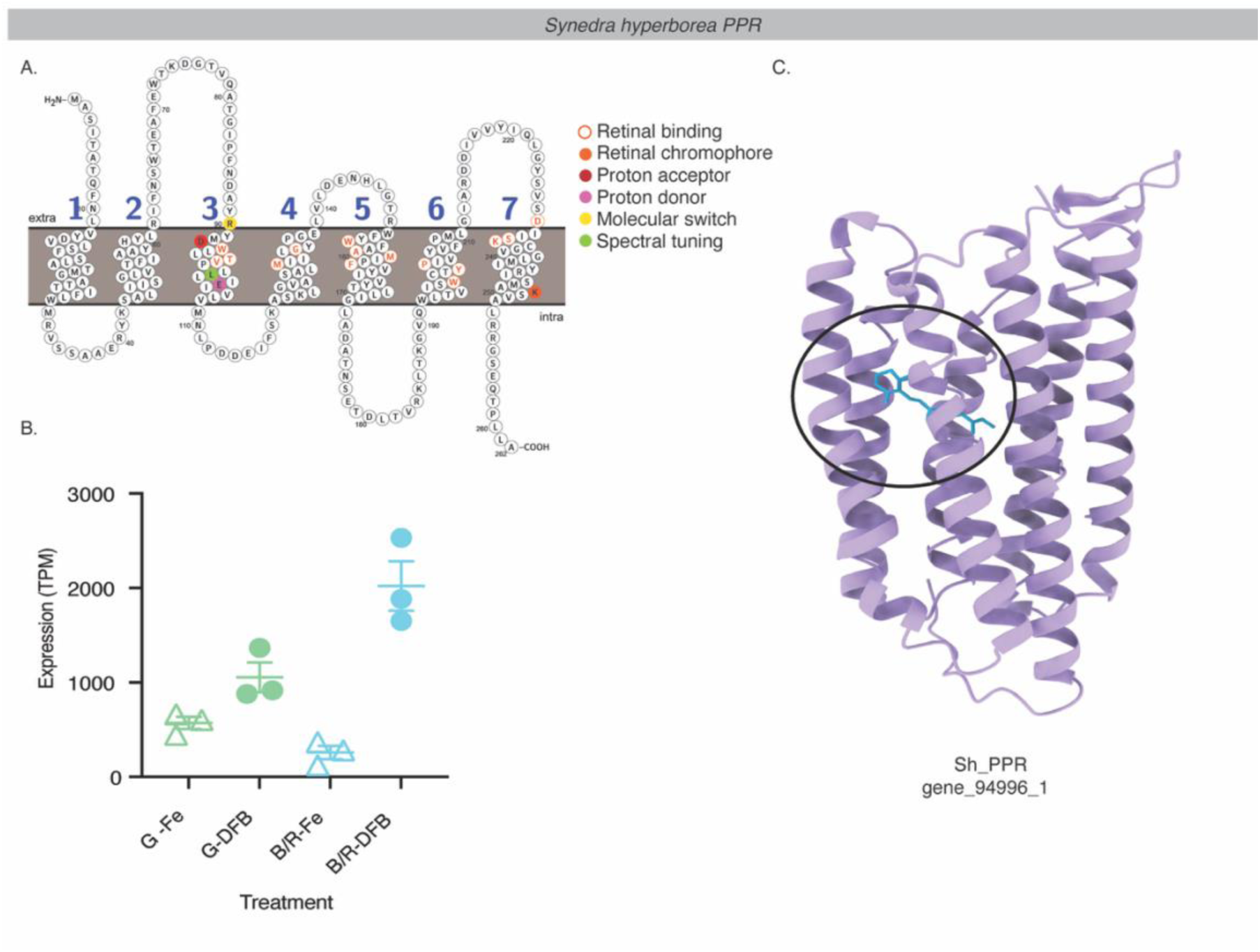
Protein structure and gene expression of dual-tuned PPR of *S. hyperborea* (Sh_PPR). (A) Two-dimensional topography of *S. hyperborea* PPR reported previously (Sh_PPR; (gene_94996_1) (23). (B) Normalized expression (transcripts per million; TPM) of Sh_PPR under varying iron conditions and light spectra. High iron and low iron replicates are shown as open triangles and solid circles, respectively; green-light treatments (G) are shown in green and blue/red-light treatments (B/R) in blue. (C) Predicted three-dimensional structure of Sh_PPR. The retinal β-ionone ring position within the fenestration zone is circled.

## Discussion

### PPR diversity underscores distinct evolutionary strategies for light acquisition

Prior studies have suggested diatoms possess a single, transcriptionally-active PPR preferentially expressed under green light and low iron availability (23, 24, 28). Consequently, acquisition of a PPR is suggested to represent a common evolutionary strategy which allows diatoms to access this narrow ecological niche. In this study, we demonstrate *P. subcurvata* contains two PPRs – one expressed highest in green light under low-iron (Ps_PPR2) – aligning with prior predictions (27), and another expressed in blue/red light under low-iron availability (Ps_PPR2). In tandem with the discovery of two distinct PPRs which are transcriptionally tuned to different light spectra, we found *S. hyperborea* contains a single PPR expressed under both light spectra during low-iron availability. These findings reshape our current understanding of the light conditions which select for PPR phototrophy in diatoms – expanding the realized rhodopsin niche. Together, these data represent distinct adaptations of light utilization within SO diatoms by underscoring a specialist vs. generalist evolutionary approach.

*P. subcurvata’s* acquisition of two PPRs transcriptionally tuned to different light spectra is consistent with a specialist strategy, where expression is tuned to a narrow niche and selected for by stable conditions (38). In contrast, *S. hyperborea’s* possession of one PPR expressed under both light spectra reflects a generalist strategy, where transcription is largely constitutive across various light niches and selected for by variable environments (39–41). Notably, these distinct evolutionary approaches to a singular gene scale to bulk cell-wide transcription. Under iron stress, light spectra elicited transcriptional reprogramming in *P. subcurvata*, with ∼50% of genes differentially expressed, whereas *S. hyperborea* transcription was largely stable. This divergence is further mirrored at the physiological level, as photopigment profiles exhibited marked shifts with light spectra in *P. subcurvata* yet remained unchanged in *S. hyperborea,* apart from the xanthophyll diatoxanthin. Collectively, we present evidence across multiple biological scales that two distinct diatoms enact different strategies to light utilization within the iron-limited SO – as underscored by this novel PPR diversity.

Yet, what distinct fitness benefits do these two diverse approaches to PPRs confer? While the adoption of a generalist PPR serves to increase tolerance of environmental variability and decrease cellular costs within the dynamic SO (42, 43) – a cost is inherent with the acquisition of two specialized, transcriptionally-tuned PPRs. We suggest this strategy may be supported by the localization of these PPRs to different organelles (Figure 6A) – which would allow for the spatial partitioning of active transport and/or ATP synthesis as a function of light spectra. In *P. subcurvata*, prior immunofluorescence investigations demonstrated Ps_PPR1 is localized to a vacuole (23), yet the localization of Ps_PPR2 remains unresolved. Based upon the literature consensus, we hypothesize Ps_PPR2 is localized a vacuole (23, 27) or the plastid (24, 44). In support of vacuolar localization (hypothesis_1_), we demonstrate Ps_PPR2 expression was coordinated with vacuolar ATPases and inorganic pyrophosphatase (Figure 6A, B) (Supplemental Table 1O), lending support for a role in light-driven ATP-synthesis within an acidic vacuole (27, 45). In contrast, a variety of genes encoding for PSII proteins were also coordinated in expression with Ps_PPR2, supporting a role in leveraging photosynthesis-driven ATP production within the plastid (hypothesis_2_). Moving forward, investigation of these hypotheses will require spatial proteomics and immunofluorescence to determine the subcellular localization of Ps_PPR2 – and additional work is required to deduce the contents of the acidified vacuole localized to Ps_PPR1 (27).

**Figure 6:**
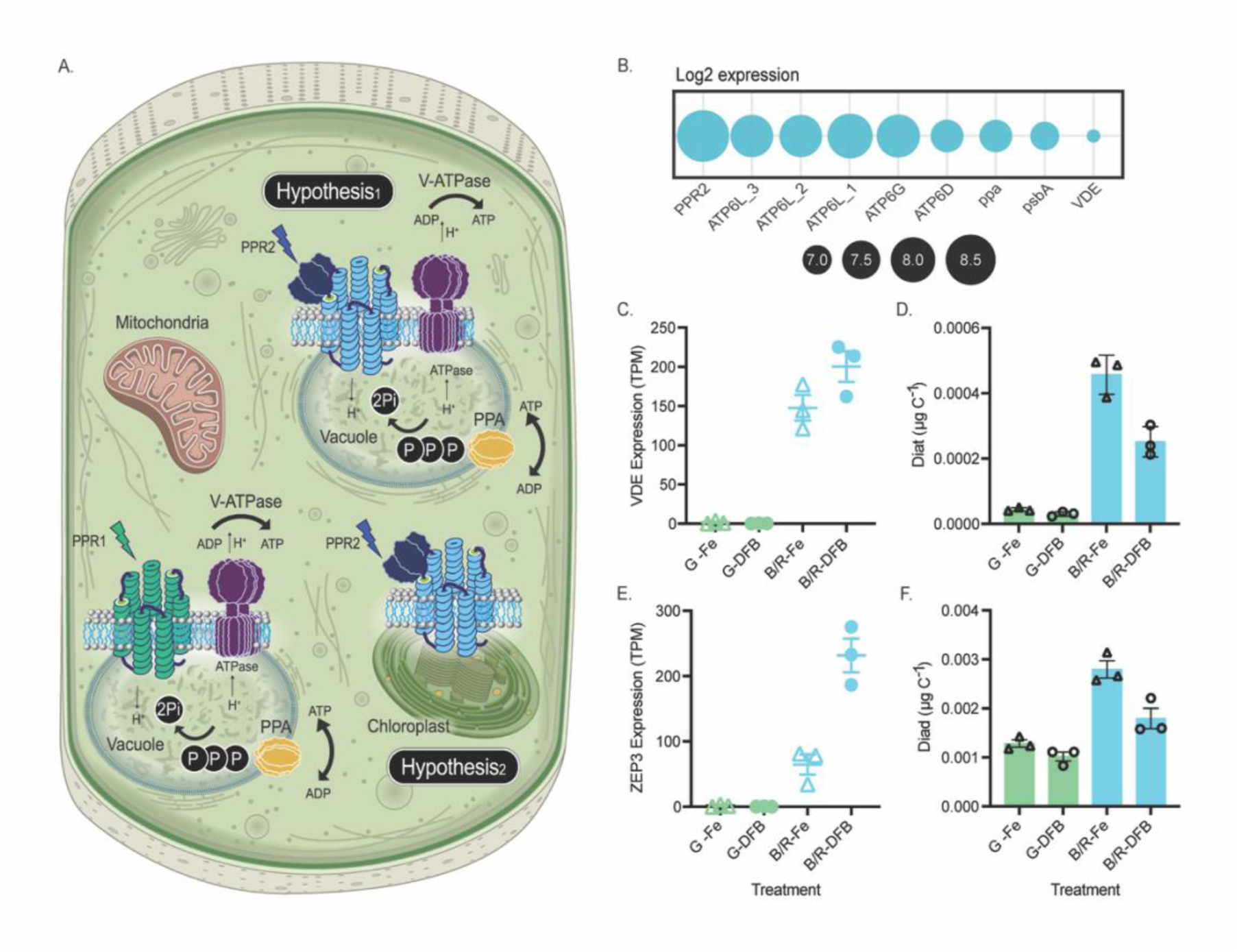
Proposed model of *P. subcurvata’s* transcriptionally tuned PPRs and the role of diadinoxanthin (Diad) and diatoxanthin (Diat) in spectral tuning to blue/red light. (A) Cellular model depicting hypotheses of spatial localization of PPRs, with Ps_PPR2 depicted as a blue-light absorbing xanthorhodopsin localized to a vacuole (hypothesis_1_) or the plastid (hypothesis_2_). (B) Log-normalized increases in mean expression of SIMPER genes of interest within low iron blue/red vs. green light treatments. Genes are all increased in expression under blue/red light – with bubble size indicating Log2 normalized expression. (C) Normalized expression of violaxanthin de-epoxidase (VDE) under varying iron conditions and light spectra. High iron (Fe) and low iron (DFB) replicates are shown as open triangles and solid circles, respectively; green-light treatments (G) are shown in green and blue/red-light treatments (B/R) in blue. (D) Concentration of diatinoxanthin in *P. subcurvata* cells normalized to cellular carbon content (µg C^-1^). For all photopigment reports, unit were calculated as follows: µg X / µg C^-1^; µg C^-1^). Bars indicate treatment means and black error bars denote standard error of the mean (SEM). (E) Normalized expression of zeaxanthin epoxidase (ZEP3) under varying iron conditions and light spectra. High iron (Fe) and low iron (DFB) replicates are shown as open triangles and solid circles, respectively; green-light treatments (G) are shown in green and blue/red-light treatments (B/R) in blue. (F) Concentration of diadinoxanthin in *P. subcurvata* cells normalized to cellular carbon content (µg C^-1^). Bars indicate treatment means and black error bars denote standard error of the mean (SEM).

### Novel expansion of the rhodopsin light niche through evidence of xanthorhodopsins

Rather than conferring fitness advantages specifically under green wavelengths of light during iron scarcity, we demonstrate two diatom PPRs are transcribed under blue/red light – with expression levels roughly two- to fourfold higher compared to green light qualities under low iron availability. While this finding expands the light niche of diatom PPRs, it contradicts the spectral paradigms of prokaryotic bacteriorhodopsins – which ascertain a leucine (Leu) at position 105 indicates the green-absorbing phenotype whereas a glutamine (Gln) determines the blue-absorbing phenotype (29). We confirmed the presence of a leucine at the spectral tuning position in all three putative diatom PPRs in this study, yet observed three distinct expression patterns as a function of light spectra – of which only one adhered to the paradigm (Ps_PPR1). While this finding indicates the prokaryotic hallmarks of spectral tuning may not be extrapolated to all eukaryotes, it also begs the question which molecular underpinnings drive spectral tuning in eukaryotes.

Explanation for the spectral tuning distinctions between *P. subcurvata’s* PPRs may lie in the ∼31 discrete substitutions present within Ps_PPR2 – which are posed to shift the hydrophobicity of this region housing the retinal chromophore. Alternatively, the blue/red spectral tuning observed in this study may lend support for the classification of diatom PPRs as xanthorhodopsins (XRs). Prior studies demonstrate dinoflagellate and diatom PPRs are phylogenetically similar to XRs (22, 24, 28, 46) – with a recent review identifying eukaryotic PPRs as “XR-homologs” (47) and another referring to diatom rhodopsins as XRs directly (24). First reported in the bacterium *Salinibacter ruber* (*48*), XRs possess an external blue light-harvesting carotenoid antenna that transfers energy to the retinal chromophore *via* a fenestration – which exposes the retinal binding pocket (49). In this study, we confirmed the presence of a fenestration in all three PPRs, yet, sequence alignments revealed tryptophan at the conserved glycine residue (G156 in *S. ruber*) (Supplemental Figure 4) – the hallmark predictor of XRs within prokaryotes (49–53). While a bulky tryptophan is thought to preclude the attachment of a xanthophyll antenna, this trait is conserved across eukaryotic XR homologs (22, 46, 54, 55) – which is likely explained by the retinal binding classification of this residue. Though further X-ray crystallography investigation is warranted, we provide novel evidence suggesting the diatom PPRs within this study are XRs. In turn, this again suggests prokaryotic hallmark predictors of rhodopsins must be applied cautiously to eukaryotic protists. Yet, the question remains – if the study proteins are XRs, what pigment is responsible for the absorption of blue light?

Across prokaryotic lineages, XRs utilize a hydroxylated carotenoid as a secondary chromophore to absorb blue wavelengths (48, 53, 56). *S. ruber* contains a salinoxanthin antenna (48), the cyanobacterium *Gloeobacter violaceus* possesses an echinenone antenna (56) and the Antarctic cyanobacterium *Pseudanabaenaceae* spp. possesses a lutein antenna (57). Yet, these pigments are conserved amongst prokaryotes – precluding their involvement in the light-harvesting antenna of a diatom XR. In this study, diadinoxanthin and diatoxanthin (DD, DE) – hydroxylated carotenoids with absorption maxima of ∼445-450 nm *λ* (blue light) (58, 59) – were the only photopigments increased in abundance under blue/red wavelengths across both diatom species. These pigments have been previously identified as components of the light harvesting antennae in diatoms (*i.e.,* the DD-DT cycle) (60, 61) and play a role in the adaptation of diatoms to light intensity *via* their interconversion (32–34). In particular, DD is largely recognized as an accessory light harvesting pigment – rendering it the ideal chromophore candidate – while DT is documented as a photoprotective, energy-dissipation pigment (62, 63). To investigate the potential for a DD light harvesting antenna of XR, we focused on comparisons within the diatom *P. subcurvata* – given the unique opportunity to compare two spectrally tuned PPRs within the same diatom.

Indeed, the expression of Ps_PPR2 increased coincidingly with violaxanthin de-epoxidase (VDE) (Figure 6B, C), which converts DD to DT (64) – and concentrations of the DT pigment were elevated in blue/red light relative to green light (Figure 6D). In tandem, we recovered a putative zeaxanthin epoxidase (ZEP3; gene_85222_3) which converts DT to DD (65, 66) and noted its expression coincided with Ps_PPR2 expression (Figure 6E) – and concentrations of the DD pigment were elevated in blue/red light (Figure 6F). Collectively, structural, transcriptional and biochemical observations support the hypothesis that Ps_PPR2 functions as an XR which may use DD as the blue-light harvesting chromophore. While this hypothesis provides a candidate pigment for diatom XRs, further investigation is required to elucidate the secondary chromophore(s) of diatom (and eukaryotic) XRs broadly.

### Spectrally tuned PPRs are posed to enhance fitness in the future polar oceans

Collectively, our findings suggest variations in light spectra fundamentally constrain the utilization of PPRs within present-day SO diatoms. In turn, this raises the question of how diatom PPRs will function within the future water column – where light climate is substantially altered. Indeed, projected increases in nutrient delivery are predicted to increase primary productivity in the Southern Ocean (16, 67, 68) – shifting the dominant wavelengths of light available in the photic zone with increasing phytoplankton biomass. In tandem, changes to ice melt are posed to reshape mixing regimes and water column structure (69–71) – altering the vertical distribution of diatoms and the dominant light spectra of the water column. Together, these factors serve to synergistically reshape the light environment of the SO, rendering spectrally tuned rhodopsins of enhanced importance in future scenarios. We subsequently explore how these changes may serve to select for spectrally tuned PPRs in the future water column.

Climate models project that intensified ice melt and altered mixing regimes will increase nutrient availability and thus primary production. In turn, these changes in phytoplankton abundance, chromophoric dissolved organic matter and suspended particles will alter the dominant light spectra of the water column due to wavelength-dependent light scattering and absorption (72–74). Indeed, nutrient-poor, oligotrophic systems are dominated by shorter, blue wavelengths of light whereas nutrient-rich, eutrophic systems are enriched in longer, red-green wavelengths (75). As a result, we hypothesize increases in SO productivity will select for PPRs spectrally tuned to green light spectra. In support, a growing body of literature indicates trophic status is a primary determinant of spectral tuning within prokaryotic rhodopsins (*i.e.,* bacteriorhodopsins) – with green-tuned rhodopsins selected for within nutrient-rich, eutrophic waters whereas blue-tuned rhodopsins are selected for in nutrient-poor, oligotrophic systems (76, 77). Building upon this, changes in biomass throughout bloom progression may exacerbate the aforementioned trends – implicating a selection for blue-tuned rhodopsins during bloom initiation and green-tuned rhodopsins during bloom maxima. Collectively, we propose these selective pressures may act upon eukaryotic PPRs – though further queries of eukaryotic rhodopsins are required within an *in situ* context.

Coincidingly, the Southern Ocean is undergoing an irreversible retreat of Antarctic sea-ice (78), which further serves to alter the light climate. While the ice-covered marine water column has been suggested to be enriched in green wavelengths of light (79), thus favoring green-tuned PPRs, a recent study demonstrated the Arctic Ocean is becoming “bluer” as sea-ice cover declines – leading to a pronounced spectral shift to shorter wavelengths in the euphotic zone (31). In turn, this study predicted spectral shifts will induce substantial changes in both the pigment and species composition of polar primary producers. Indeed, it was recently proposed that blue-absorbing XRs play a role in Antarctic *Pseudanabaenaceae*’s acclimation to prolonged low light intensities and spectral shifts in the Southern Ocean (57) – which lends further support to the current study which was conducted under low-light conditions (*i.e.,* 10 µmol photons m^-2^ s^-1^) in effort to simulate the co-limitation of light intensity and iron availability. Cumulatively, our findings extend this growing importance of blue-tuned rhodopsins to eukaryotic phytoplankton (*e.g.,* diatoms) which routinely dominate the Southern Ocean.

Combining our current results with previous findings, we suggest future increases in productivity will select for green-tuned diatom PPRs within the Southern Ocean – whereas decreasing ice cover will select for blue-tuned diatom PPRs. More broadly, we hypothesize that seasonal bloom progression serves to further shape PPR selection – with bloom initiation (bluer waters and lower biomass) selecting for blue-tuned PPRs and bloom maxima (greener waters and higher biomass) selecting for green-tuned PPRs. Considering these scenarios are likely to coincide, further investigation into the relative contributions of each driver will be necessary to discern competitive trade-offs and intraspecific competition. In turn, this uncertainty illustrates the need for further investigation to ensure putative PPR fitness advantages can be extrapolated to large-scale biogeochemical consequences and incorporated into regional ecosystem models. Cumulatively, our findings reinforce the notion that climate change in increasingly manifesting as a “light problem” within the global oceans and suggest that spectral tuning of PPRs represents an underappreciated mechanism of adaptation which warrants attention – particularly within the context of dynamic light climates of the SO.

## Materials and Methods

### Polar diatom isolates

The polar diatoms *Pseudo-nitzschia subcurvata* (UNC1901) and *Synedra hyperborea* (UNC1904) were previously isolated from the Palmer Long-Term Ecological Research (Pal-LTER) region of the western Antarctic Peninsula and identified using morphological and molecular (18S rRNA) methods (23). The presence of a PPR in each diatom was confirmed *via* transcriptomics (80) and the cellular localization of the *P. subcurvata* PPR was determined by immunofluorescence microcopy (23).

### Growth assays under varying iron availability and light spectra

Cultures were maintained in AQUIL synthetic seawater medium supplemented with vitamins and nutrients (300 µM NO₃^-1^, 20 μM PO₄^-3^, 200 μM Si(OH)₄, and 100 μM EDTA) (81). Cells were acclimated to conditions representative of the Southern Ocean (4 °C, 10 µmol photons m^-1^ s^-2^, 24 h irradiance) under varying iron availabilities and light spectra. Iron replete (1,370 nM Fe:100 µM EDTA) or iron-limiting (4nM Fe:400nM Desferrioxamine-B (DFB)) treatments were incubated under blue/red (peak emissions at 440 & 660 nm *λ*, respectively) or green (peak emission at 530 nm *λ*) light. Light treatments were established using custom engineered light banks (Supplemental Figure 5). Photon flux densities of the specific high power single wavelength LEDs were tuned using Arduino IDE software (v.1.6.4+) pulse width modulation. Spectral emission was verified by UV-Vis Spectrometry (OceanOptics, Orlando, FL, USA) (Supplemental Figure 6). Following acclimation (∼6 weeks of three sequential transfers), 1 L cultures were inoculated in biological triplicate (n = 3) under trace-metal clean conditions. Growth was monitored by *in vivo* Chl *a* autofluorescence using a Turner Designs model 10-AU fluorometer (82). Specific growth rates were calculated as the linear regressions of the natural log of Chl *a* autofluorescence vs. time. Samples were harvested during late exponential growth for subsequent analyses.

### Cellular C:N ratios

Cellular carbon and nitrogen ratios (C:N) were determined as described prior (83). Briefly, 50 mL subsamples were filtered onto pre-combusted GF-75 filters (450 °C for 6 h) (Advantec MFS, Dublin, CA, USA), flash frozen in liquid nitrogen and stored at -20 °C. Prior to analysis, filters (n = 24) were dried at 65 °C for 24 h and encapsulated in tin. Total cellular carbon and nitrogen (μg) were quantified using the Costech 4010 CHNOW elemental combustion system and U.S. EPA method 440.0 (84). Three blanks were run in tandem and confirmed to be below the limits of detection (0.071 mg C and 0.005 mg N). Samples were not acidified and thus included cellular inorganic carbon.

### Photophysiology and photopigment analyses

Photophysiological measurements were conducted as previously described (23). Briefly, 10 mL subsamples were acclimated to low light (∼10 μmol photons m^-2^ s^-1^) for 30 minutes and analyzed using a Fluorescence Induction and Relaxation (FIRe) fluorometer (Satlantic). Single-turnover flashes were applied to PSII reaction centers for 50 iterations (85), followed by an 80 ms saturating pulse of blue light (450 nm *λ*). Maximum photochemical efficiency of PSII (Fv•Fm^-1^) was calculated using the standard FIRe induction-relaxation protocol.

Cellular photopigments were quantified as previously described (86). Briefly, 100 mL subsamples were filtered onto GF-75 filters, flash-frozen in liquid nitrogen, and stored at −80 °C until processing at the *University of South Carolina Estuarine Ecology Laboratory.* Samples (n = 24) were freeze dried and extracted in 90% acetone at -20 °C for 24 h (87). Extracts were processed *via* high-performance liquid chromatography using 22 primary algal pigment standards (88–90). Total photopigment concentrations (µg) were normalized to total cellular carbon.

### RNA extraction and sequencing

For RNA extractions, 500 mL subsamples were filtered onto 5.0 μm nominal pore size 47-mm diameter polycarbonate filters, flash frozen in liquid nitrogen and stored at - 80 °C. Filters were bead-beaten in lysis buffer, and total RNA was extracted using the RNAqueous^TM^-4PCR Total RNA Isolation Kit (Invitrogen, Carlsbad, CA, USA). Residual DNA was digested using the TURBO DNA-free Kit (Invitrogen). RNA concentrations were quantified using the Qubit RNA HS Assay Kit (Invitrogen). PolyA-selected libraries were prepared and sequenced by Azenta Life Sciences (Indianapolis, IN, USA) on the Illumina HiSeq platform to yield ∼350 million, 150 bp, paired-end reads.

### Transcriptome assembly and annotation

Bioinformatic analysis was performed using an established transcriptomic workflow (91, 92). Raw libraries (n = 24) were quality assessed using FastQC (v.0.12.1). Adapter trimming and quality filtering were performed with Trimmomatic (leading = 20, trailing = 20, sliding window = 4:15, minimum length = 36) (v.0.39) (93). Paired reads were interleaved using BBTools and residual rRNA was removed with BBMap (v.38.18) (94). Libraries of *P. subcurvata* (n = 12) and *S. hyperborea* (n = 12) were concatenated and assembled (coassembled) into species-specific coassemblies using MEGAHIT (v.1.2.9) (95). Assembly quality was assessed with QUAST (v.5.2.0) (96) and reads were mapped back to each respective coassembly with BBMap. Open reading frames (genes) were predicted with MetaGeneMark2 (v.1.0) (97). Functional annotation was performed with eggNOG using a specified e-value of < 1e^-10^ (v.2.1.12) (98) and the standard reference database (v.5.0.2). Given diatom isolates were not axenic, taxonomic annotation was performed with EUKulele (v.1.0.6) (99) using the PhyloDB reference database (v.1.076). Read counts to predicted genes were tabulated with featureCounts within the subread package (v.2.0.1) (100). Only genes assigned as eukaryotic were retained for downstream analyses, prokaryotic sequences attributed to the presence of heterotrophic bacteria were omitted.

### Identification and structural characterization of PPRs

Given PPRs remain poorly resolved and underrepresented in reference databases (101) (92), candidate genes initially annotated as bacteriorhodopsins by eggNOG were subjected to additional validation. Protein sequences were queried against the NCBI nonredundant database to verify the top hit, based on e-value and sequence identity, corresponded to a previously-described diatom PPR. Two dimensional membrane topologies were predicted with Protter (v.1.0) (102) to confirm hallmark characteristics of rhodopsins and conserved functional domains were identified using the Simple Modular Architecture Research Tool (v.10.0) (103). Three-dimensional structures were predicted by AlphaFold 3 (104) and aligned by the retinal β-ionine ring position in UCSF ChimeraX (v.1.11.1) (105) and examined for fenestrations as reported prior (36).

### Statistical analyses

Statistical analysis of physiological data was performed using two-way ANOVA post-hoc multiple comparisons (Tukey’s HSD) in GraphPad Prism (v.10.3.0). Statistical analysis of molecular data was performed in R (v.4.2.2). Normalization of raw reads to transcripts per million (TPM), calculation of Bray-Curtis dissimilarity and nonmetric multidimensional scaling (nMDS), and estimatation of Similarity Percentage (SIMPER) between libraries was performed using customized R scripts as reported prior (92). Differential Expression (DE) analysis was performed using DESeq2 (v.1.44.0) (106) to determine statically significant trends (padj < 0.05, abs Log2FC >1). Pairwise comparisons were performed between treatments with subsequent overlaps deduced between all four treatments.

## Acknowledgments

We thank Raquel Flynn, Margarita Lankford and Alexander Truchon for their constructive conversations and assistance with the interpretation of this data. We thank Melissa Smith of the UNCW Center for Marine Science for her collaboration and scientific illustrations We are grateful to the captain and crew of the *ARSV Lawrence M. Gould* and participants of the Pal-LTER 2018 cruise for their assistance in the collection of diatom isolates. Portions of this work were performed on the *UNC Chapel Hill High Performance Computing Cluster Longleaf* and the *UNCW High Performance Computing Cluster Cuttlefish*. This work was funded by an NSF Oceans Science Postdoctoral Research Fellowship OCE-2307229 (BNZ) and NSF OPP-1745036 (AM). The presentation of this work at the 2025 Gordon Research Conference in Polar Marine Science was made possible by funds provided through the AGU Leadership Academy and Network for Diversity and Inclusion in the Geosciences (AGU LANDInG) Program (BNZ).

## Data availability

The raw transcriptomes (n = 24) of this study are publicly available on the National Center for Biotechnology Information Sequence Read Archive (NCBI SRA) under [BioProject PRJNA476717](https://www.ncbi.nlm.nih.gov/bioproject/PRJNA1476717). A complete list of the BioSample and SRA accessions is provided in the supplemental file of this publication. The concatenated assembly and open reading frame protein files are publicly available on the lead author’s [Zenodo](https://zenodo.org/records/20869403). Biochemical and growth data are available in the supplemental file of this publication. Data is available upon request.

