## Supplementary material for "Spectral tuning of proton-pumping rhodopsins underscores distinct light adaptations in polar diatoms": ZEPERNICK ET AL 2027 SUPPLEMENTAL SUBMITTED PNAS.pdf

[Φ]Current affiliation: Centre of Excellence Node Administrator, Research School of Chemistry, Australian National University, Canberra, Australia
[#]Current affiliation: School of Environmental Sciences, University of East Anglia, Norwich, United Kingdom
[£]Current affiliation: School of Earth and Ocean Sciences and Department of Biology, University of Victoria, British Columbia, Canada

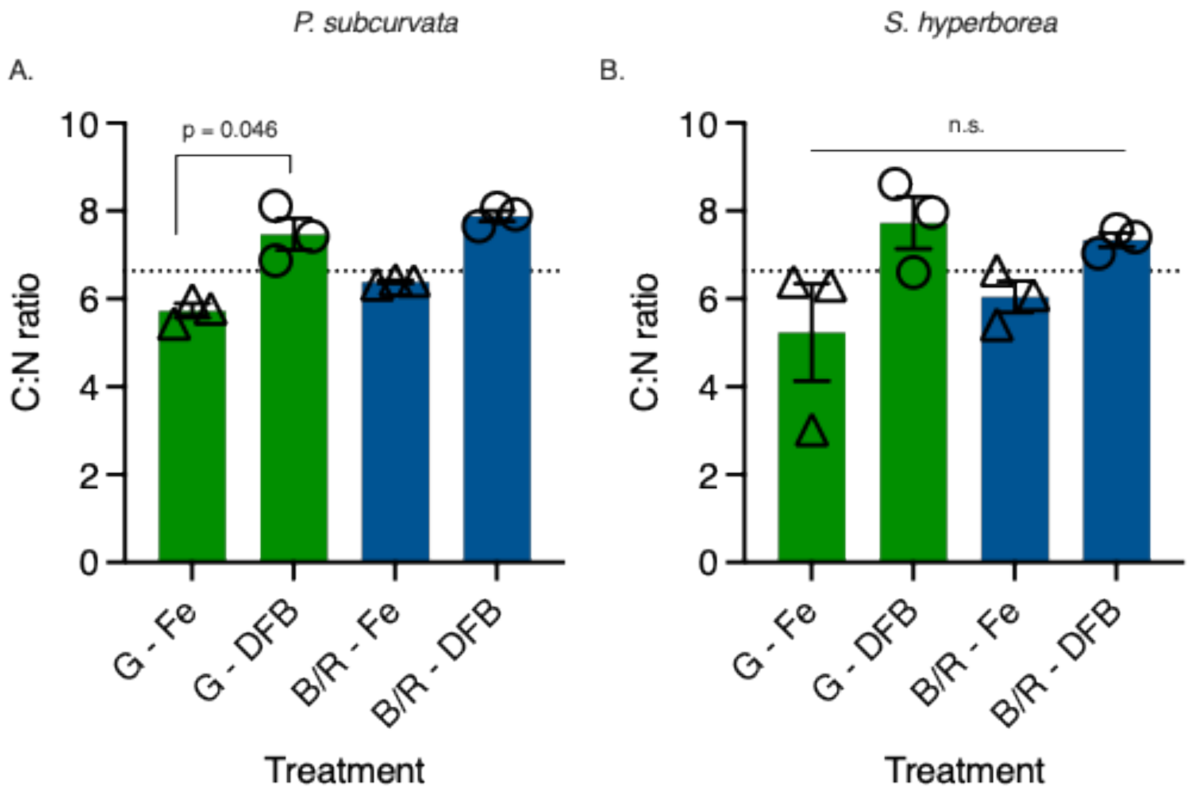


**Supplemental Figure 1:** Cellular C:N ratios of *P. subcurvata* (A) and *S. hyperborea* (B) under experimental iron and light treatments. Green light treatments are indicated by dark green bars whereas the blue/red light treatments are depicted by dark blue bars. High iron (1370 nM Fe) treatments are depicted by triangles and low iron (DFB) treatments are depicted by circles. The theoretical optimal C:N ratio of phytoplankton cells based on the Redfield Ratio (6.6) is depicted by a dotted black line. Statistically significant ($p < 0.05$) differences between treatments of interest (high iron vs. low iron and green light vs. blue/red light) are indicated where applicable. Otherwise, n.s. = not significantly different across treatments.

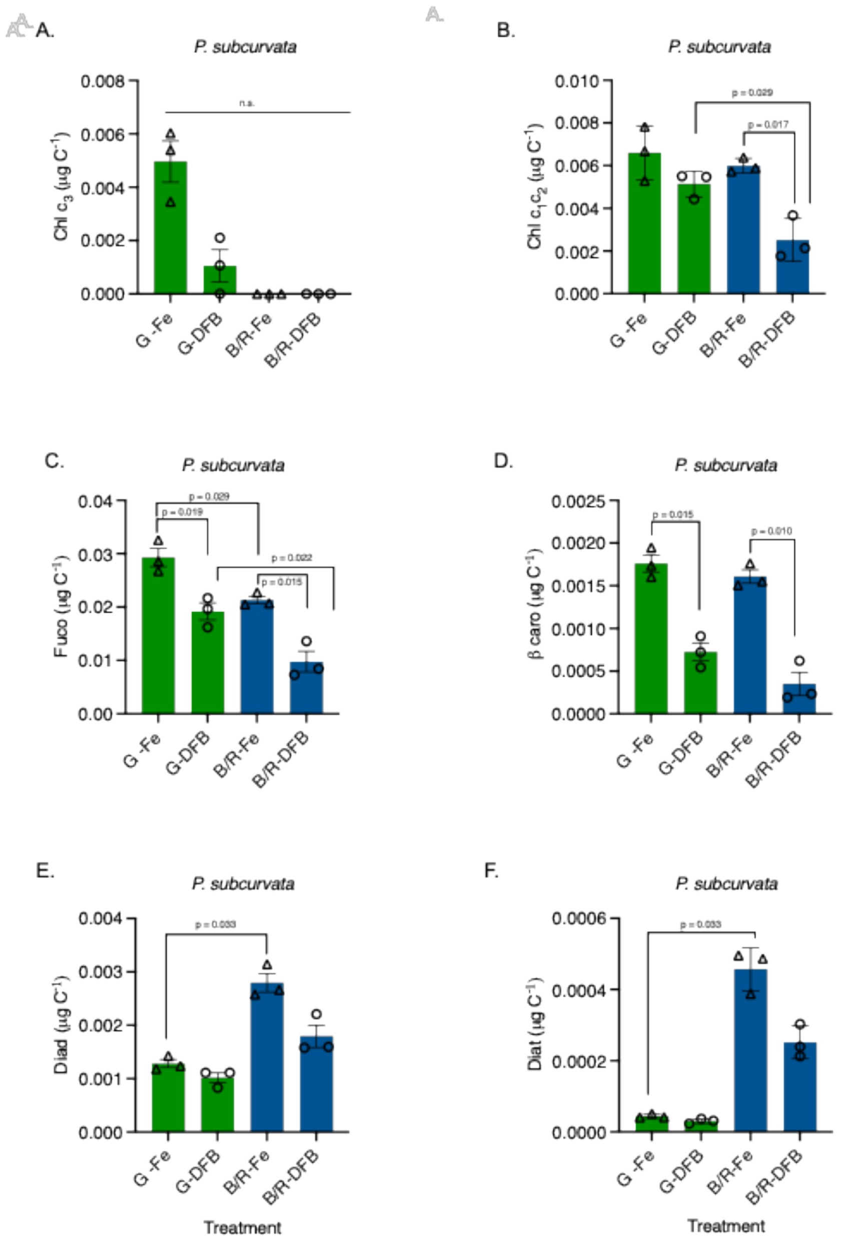


**Supplemental Figure 2:** *P. subcurvata* cellular photopigment concentrations normalized to cellular carbon content (µg $C^{-1}$). In this case, both cellular photopigment and carbon concentrations were initially reported in µg prior to normalization and are thus defined as µg pigment per µg cellular carbon in µg $g^{-1}$. (A) Chlorophyll $c_3$, (B) Chlorophyll $c_1c_2$, (C) Fucoxanthin, (D) ß-carotene, (E) Diadinoxanthin, (F) Diatoxanthin. Green light treatments are indicated by dark green bars whereas the blue/red light treatments are depicted by dark blue bars. High iron (1370 nM Fe) treatments are depicted by triangles and low iron (DFB) treatments are depicted by circles. Statistically significant ($p < 0.05$) differences between treatments of interest (high iron vs. low iron and green light vs. blue/red light) are indicated where applicable. Otherwise, n.s. = not significant across treatments of interest. Chlorophyll $c_3$ was not detected in *P. subcurvata* blue/red light treatments.

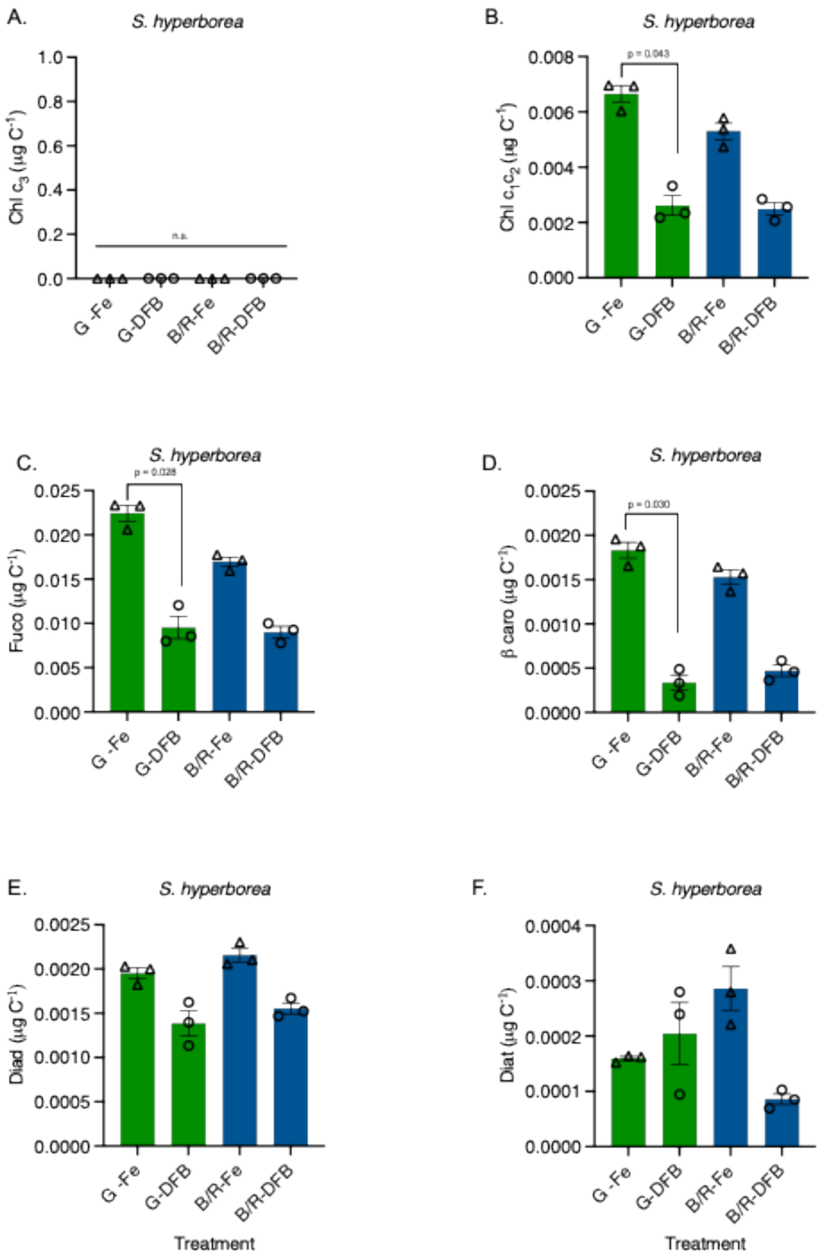


**Supplemental Figure 3:** *S. hyperborea* cellular photopigment concentrations normalized to cellular carbon content ($\mu g\ C^{-1}$). In this case, both cellular photopigment and carbon concentrations were initially reported in µg prior to normalization and are thus defined as µg pigment per µg cellular carbon in $\mu g\ g^{-1}$. (A) Chlorophyll $c_3$, (B) Chlorophyll $c_1c_2$, (C) Fucoxanthin, (D) ß-carotene, (E) Diadinoxanthin, (F) Diatoxanthin. Green light treatments are indicated by dark green bars whereas the blue/red light treatments are depicted by dark blue bars. High iron (1370 nM Fe) treatments are depicted by triangles and low iron (DFB) treatments are depicted by circles. Statistically significant ($p < 0.05$) differences between treatments of interest (high iron vs. low iron and green light vs. blue/red light) are indicated where applicable. Otherwise, n.s. = not significant across treatments of interest. Chlorophyll $c_3$ was not detected in any *S. hyperborea* light / iron treatments.

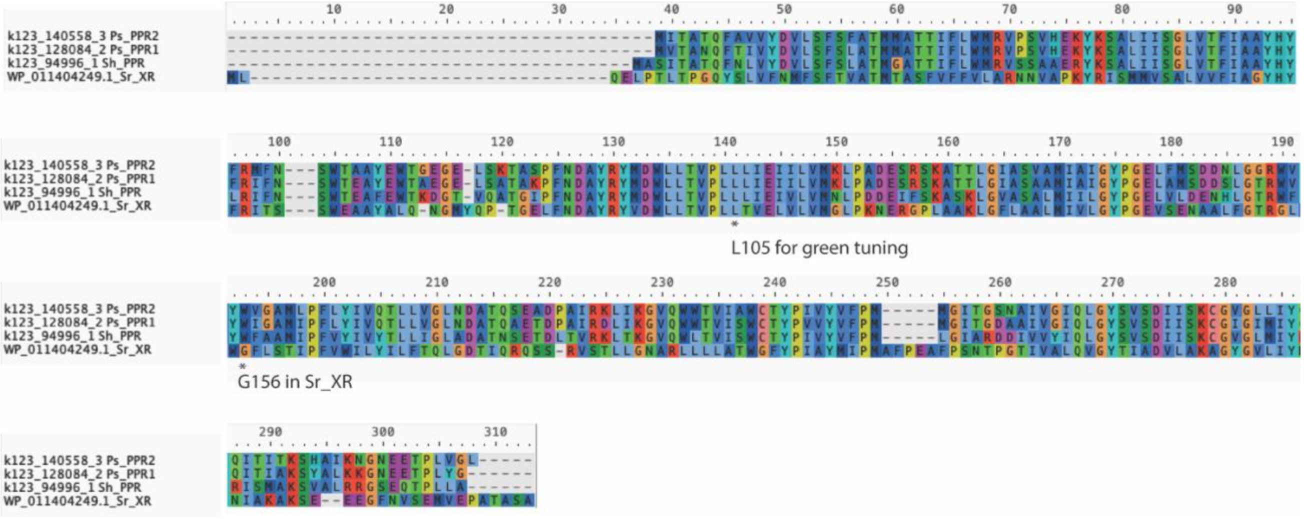


**Supplemental Figure 4:** Alignment of the three diatom PPRs recovered in this study (Ps_PPR1, Ps_PPR2 and Sh_PPR) to the xanthorhodopsin sequence originally described in *Salinobacter ruber.* Ungapped residue numbers were used to determine the location of conserved amino acid residues of interest. The conserved spectral tuning residue for green-light absorbing rhodopsins (L105) is indicated with an asterisk (*). In addition, the conserved G156 reported in XRs as the hallmark for the carotenoid-binding antenna is indicated with an asterisk (Marín et al., 2025).

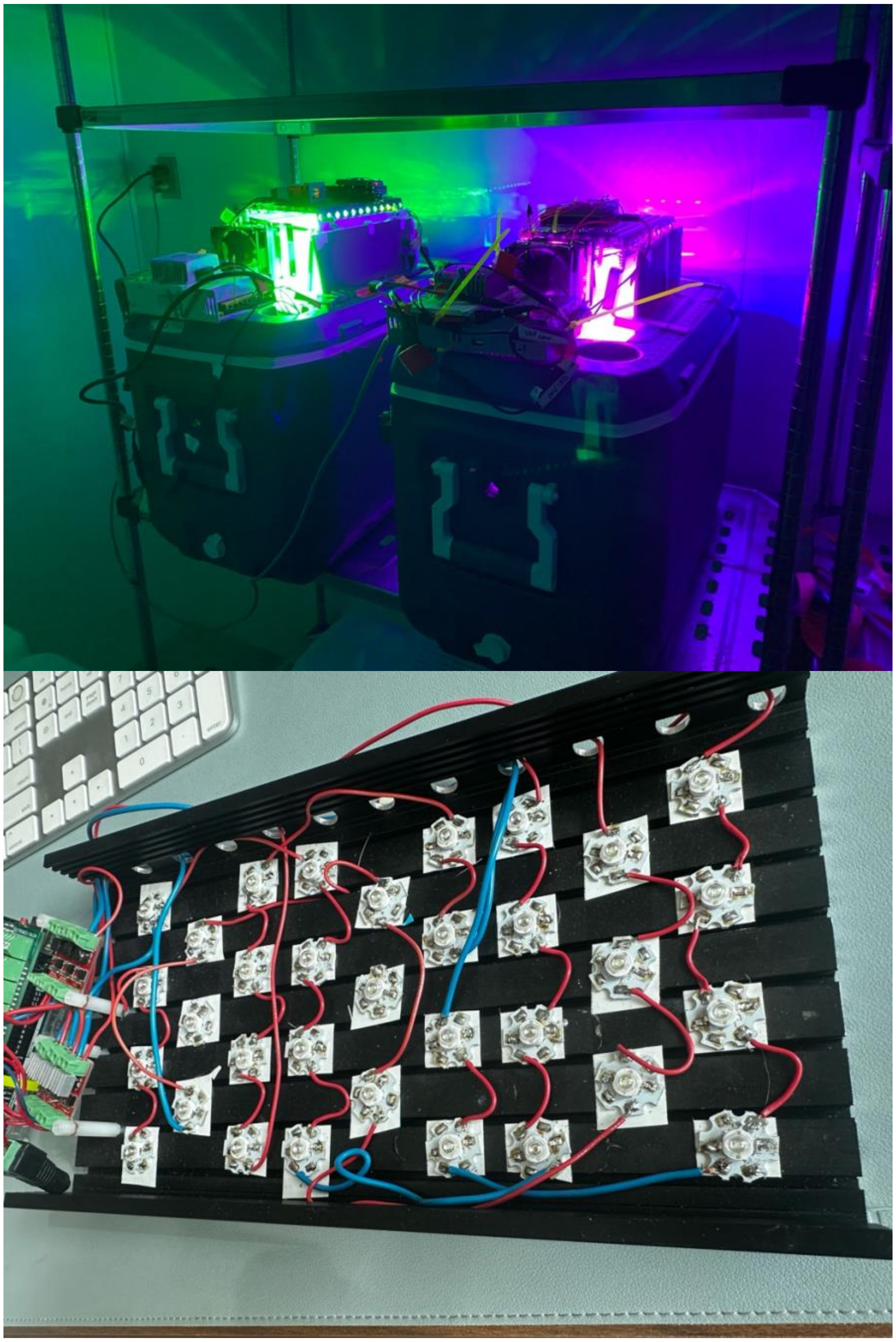

**Supplemental Figure 5:** Custom engineered light banks (A, B). High power single wavelength LEDs were tuned to photon flux density using Arduino microcontrollers and pulse width modulation.

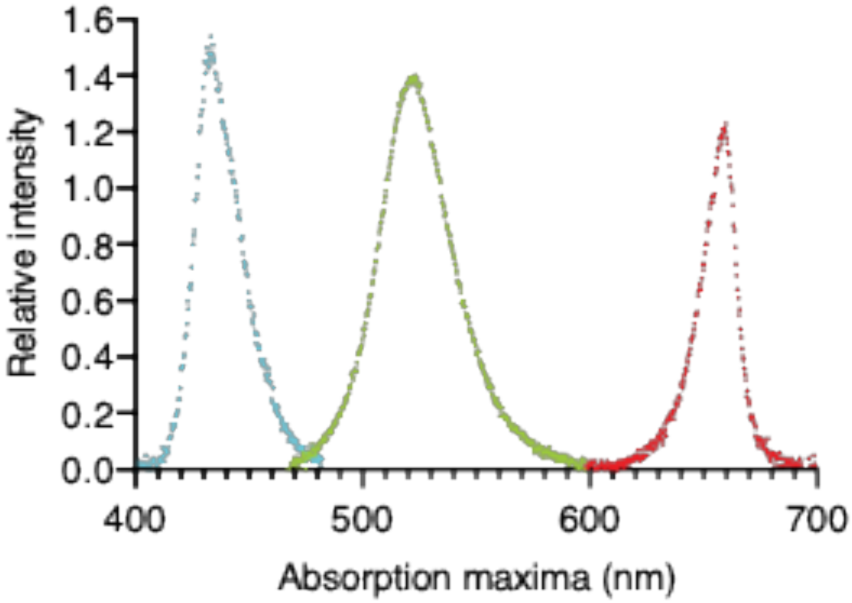


**Supplemental Figure 6:** Spectral absorption profiles of the three different LEDs used in the light banks. Spectra were measured *via* UV-VIS spectroscopy. Blue and red peaks correspond to the blue/red light incubators and green peaks correspond to the green light incubators.

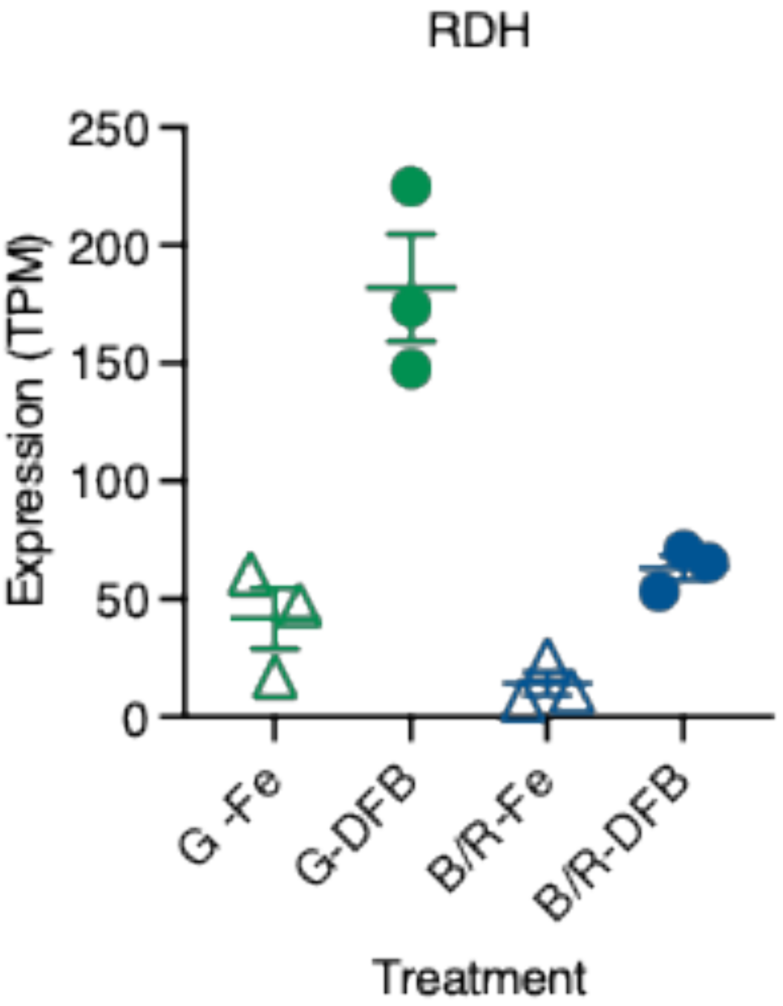


**Supplemental Figure 7:** *S. hyperborea* expression (normalized to transcripts per million; TPM) of retinal dehydrogenase (RDH) which catalyzes the oxidation of retinal into retinoic acid. Green light treatments are indicated by green shapes whereas the blue/red light treatments are depicted by dark blue shapes. High iron (1370 nM Fe) treatments are depicted by open triangles and low iron (DFB) treatments are depicted by solid circles.
